# Fine scale population structure linked to neutral divergence in the common sole (*Solea solea*), a marine fish with high dispersal capacity

**DOI:** 10.1101/662619

**Authors:** Alan Le Moan, Belén Jiménez-Mena, Dorte Bekkevold, Jakob Hemmer-Hansen

## Abstract

The Baltic Sea provides a classical example of how an environmental gradient is associated with the distribution of marine species. Here, numerous genetic studies have revealed clear patterns of population structuring linked to the physical features of the gradient itself. Nevertheless, it remains difficult to distinguish clearly between the different micro-evolutionary processes that shape these structured populations *in situ*. The common sole (*Solea solea*) is a benthic flatfish that rarely occurs within the Baltic Sea, but that exhibits a clear genetic break between populations from the North Sea – Baltic Sea transition zone and the remainder of the Atlantic Ocean. Here, we aim to evaluate the extent to which natural selection is involved in the observed patterns of divergence of sole populations occurring in the transition zone by comparing them with population structures of other flatfish species that have successfully colonized the Baltic Sea. By using several thousand of ddRAD-derived SNPs, we identified a fine-scale pattern of isolation-by-distance (IBD) of sole populations in the region. However, despite strong biological similarities among the flatfishes compared here, the sole IBD was, by far, the lowest detected across the transition zone. While selection was inferred to strongly influence all other flatfishes evolutionary histories, the analytical inference on the sole demographic history suggests that this fine-scale IBD is mainly maintained by neutral processes due to low effective population size of sole in the transition zone and asymmetrical gene flow. Our work contributes to a growing body of evidence suggesting that the strength of the different micro-evolutionary processes is species-specific, even when species occur in the same environment.

## Introduction

The study of local adaptation and ecological speciation often relies on species that have been successfully established across environmental clines. Nevertheless, comparing between successfully and unsuccessfully colonizing species occurring in the same environmental gradient can provide relevant data for improving our understanding of the ecological divergence continuum.

The Baltic Sea provides a classical example of a marine environmental gradient, ranging from near freshwater in its inner parts to brackish and nearly marine in the western parts. This area was only recently connected to the Atlantic Ocean (8 000 years ago), and it has subsequently been colonized by marine fauna (Björck, 1995). Several studies have identified clear patterns of population structure associated with the gradient (Bekkevold et al., 2005; Cuveliers et al., 2012; Hemmer-Hansen et al., 2007; Limborg et al., 2009; Nielsen *et al.*, 2004; Riginos and Cunningham, 2005; Väinölä and Hvilsom, 1991). These may have evolved through several evolutionary processes (reviewed in Johannesson and André, 2006), aided by the unique oceanographic characteristics of the region. For example, the Baltic Sea basin is connected to the North Sea through a narrow transition zone. This single point of contact is associated with asymmetric oceanographic gyres that limit the inflow from the Atlantic into the Baltic Sea (Naumann *et al.*, 2017), which in turn may limit the connectivity between populations from each side of the transition zone. Consequently, these populations are more likely to diverge by genetic drift due to this potential isolation at the margin of the species distribution (Johannesson and André, 2006). Furthermore, the transition zone itself is characterized by sharp environmental contrasts that are expected to lead to a primary zone of divergence driven by local adaptation (Nissling and Westin, 1997). Finally, such environmental gradients are expected to retain reproductive incompatibilities through the coupling between endogenous and exogenous barriers to gene flow, resulting in secondary zones of divergence (Barton, 1979; Bierne *et al.*, 2011). Hence, discrete population structure can evolve in response to these distinct processes of divergence, all resulting in genetic breaks localized along the common transition zone.

While accurately disentangling the genetic signatures of these three types of evolutionary processes has been challenging with the use of few genetic markers, it has become increasingly feasible with the access to genomic data. Such data provide the resolution to detect signatures of selection within specific regions of the genome (Hemmer-Hansen *et al.*, 2013; Nielsen *et al.*, 2012). Further, the increase in numbers of genetic markers allows us to explore recent population history events and to distinguish among neutral, primary and secondary processes of divergence (Butlin *et al.*, 2014; Le Moan *et al.*, 2016; Rougemont *et al.*, 2017, 2017). In Le Moan et al. (2019a), we found evidence of both primary and secondary zones of divergence associated with the colonization of the Baltic Sea in four flatfish species: the turbot (*Scophthalmus maximus*), the common dab (*Limanda limanda*), the European plaice (*Pleuronectes platessa*), and the European flounder (*Platichthys flesus*). Although the distributions of plaice and dab within the Baltic Sea is limited to the westernmost region, while flounder and turbot occur across most of the basin, all these species can be considered as successfully established in the Baltic Sea.

The common sole (*Solea solea*) is a benthic flatfish species that is not as well established within the Baltic Sea as the previous species, as it rarely occurs beyond the main environmental transition zone into the Baltic Sea (Storr-Paulsen *et al.*, 2012). Previous population genetic studies in the common sole shown weak structuring across the northeastern Atlantic (Kotoulas *et al.*, 1995; Rolland *et al.*, 2007), which exhibited, nonetheless, clear patterns of isolation-by-distance (IBD) along the continental shelf (Cuveliers *et al.*, 2012; Diopere *et al.*, 2018). This overall IBD suggests limited connectivity among the spawning grounds of sole throughout the northeastern Atlantic (Guinand *et al.*, 2008). Furthermore, the clearest genetic break matched the North Sea and the Baltic Sea transition zone. Here, several outlier loci where identified that could potentially reveal genomic footprints of selection, driven by local adaptation to the environmental gradient (Diopere *et al.*, 2018; Nielsen *et al.*, 2012). However, these outliers were also clearly associated with an IBD pattern, and no major genetic cline was detected between the North Sea and Baltic Sea populations. Therefore, the higher divergence levels in the transition zone could also result from isolation at the margin of the sole distribution (Johannesson and André, 2006). In order to understand the evolutionary processes behind the observed differentiation between the populations of sole in the transition zone, we aim to: 1) re-evaluate the fine-scale population structure along this area using improved genomic coverage to assess the extent to which local adaptation is involved in maintaining population structure; and 2) compare the spatial structure of sole to that of other flatfishes in the area that differ in their ability to colonize the environmental gradient. Here, we show that even in the absence of a reference genome, we can draw important inferences regarding the evolutionary consequences of population structuring in an environmental transition zone with the use of a high number of genetic markers. We find that the IBD pattern detected at large geographical scale is also evident at finer scale. Analytical inference on the demographic history suggests that the fine-scale IBD is mostly maintained by neutral processes. Although few loci show evidence for selection leading to relatively steep allelic frequency clines in the area, the extent to which selection is involved in structuring sole populations is much less important than in the other flatfish species established in the Baltic Sea. Our work contributes to a growing body of evidence suggesting that the strength of the different micro-evolutionary processes is species-specific, even when species occur in the same environment.

## Material and Methods

### Sampling

We sampled 146 common sole individuals (*Solea solea*) at the spawning season between 2016 and 2018 from six sites spread along the coast of Denmark (Figure 1A). The common sole is a relatively rare species in comparison to other flatfish in the area, and its density decreases along the North Sea – Baltic Sea transition zone. Therefore, the sample sizes varied between sites, ranging from 17 to 35 individuals per site. However, the sample sizes were sufficient to explore the main genetic structure along the transition zone.

**Figure 1:**
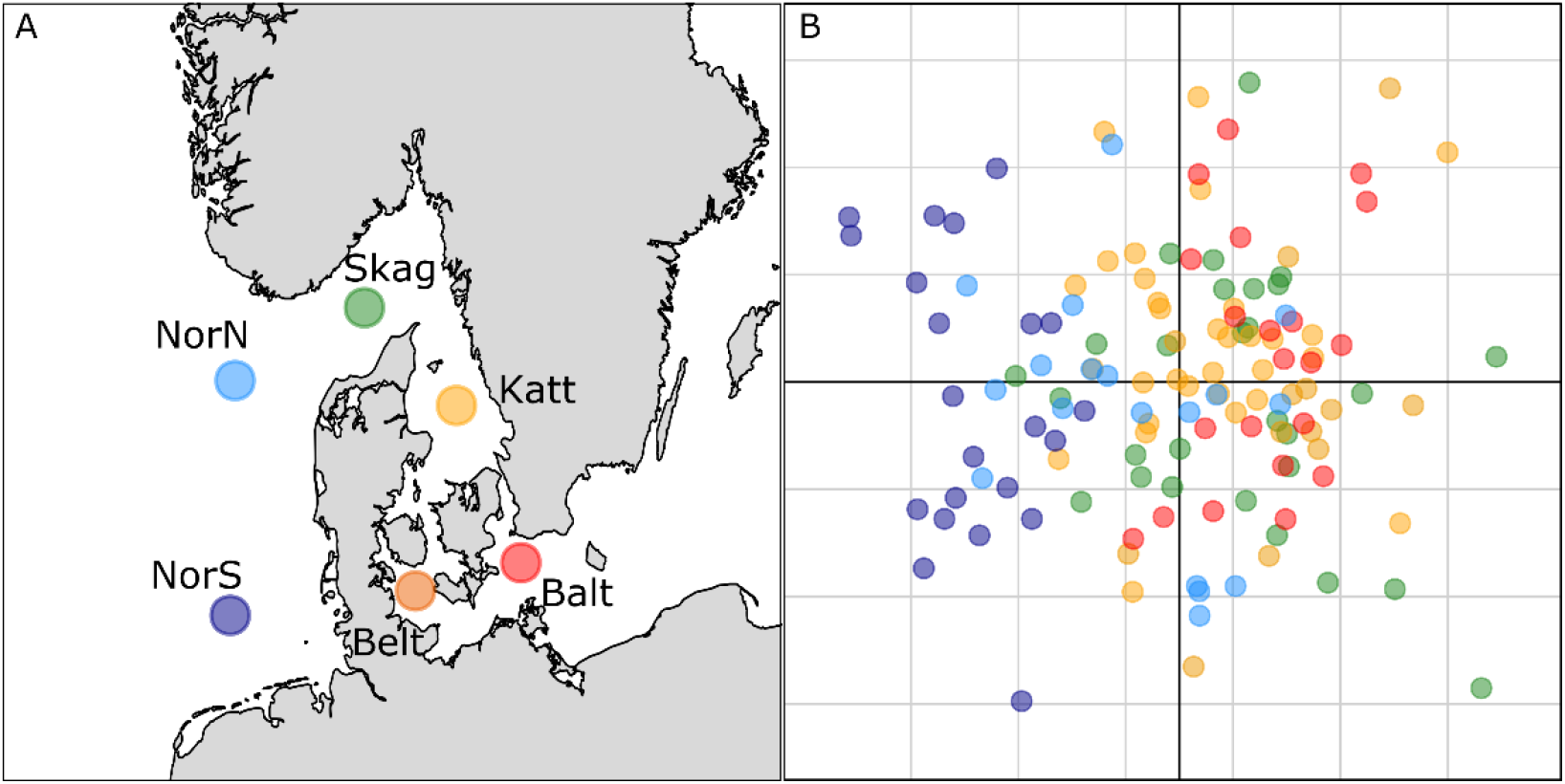
Sampling map for sole across the North Sea – Baltic Sea transition zone (A) and PCA of individual genetic diversity (B). The color on the PCA in B matches the sampling locations depicted in the map in A

### DNA extraction and preparation of double-digestion Restriction-site Associated DNA (ddRAD) libraries

Individual genomic DNA was extracted with the DNeasy Blood Tissue kit (Qiagen). Individual DNA extractions were processed with two restriction enzymes, Msp1 and Pst1, with rare and frequent cutting site, respectively. We randomly pooled 70 and 76 barcoded individual in equimolar proportion (20 ng/ul per sample) and built two independent ddRAD libraries following a modified protocol of Poland and Rife (2012). Each library was size-selected on agarose gels to retain insert sizes between 300 and 400 base pairs (bp). Following PCR amplification (12 cycles), the two libraries were purified using AMPure^®^ beads (AgenCourt). Finally, the quality of the libraries was visualized using a Bioanalyzer^®^ 2100 (Agilent Technologies), using the High Sensitivity DNA reagent kit. The two libraries were sequenced in paired-ends (2 x 100 bp) on two independent Illumina HiSeq4000 lanes.

### Bioinformatics pipeline

The pooled libraries were processed using the pipeline developed in Jiménez-Mena *et al*., in prep. The sole samples were demultiplexed by barcodes using the function process_ratags.pl from stacks 1.46 (Catchen *et al.*, 2013). Only reads with a quality of sequencing above 10 were kept (Figure S1). One of the barcodes from the second library was one bp longer than the barcodes used in the first library. Consequently, all the forward reads were trimmed first to 90 bp to be consistent across libraries. Forward and reverse reads with more than 10 overlapping base pairs were merged together using flash (Magoč and Salzberg, 2011). The merged and unmerged reads were extracted in different fasta files for each individual. Merged reads were trimmed to 171 bp and smaller reads were discarded from the analyses. For the unmerged reads, the first 13 bp and the last 6 bp were removed from the reverse reads. Using custom bash scripts, reverse reads were then concatenated with the forward reads to obtain sequences of same length (171 bp) as the merged reads. Merged and concatenated reads were then pooled together in individual fasta files. More details about this pipeline can be found in the supplementary material of Jiménez-Mena *et al*., in prep. SNPs were called using the *de novo* pipeline of Stacks 1.46 (Catchen *et al.*, 2013). Specifically, for each individual, we used the ustacks function set with a minimum coverage of five to considered a stack of identical reads as biological sequences (m = 5), and a maximum of five differences (M = 5) to consider two stacks of reads and secondary unstacked reads as homologous sequences. Then, all sequences were registered into a catalogue using the function cstacks, allowing six differences (n = 6) between the individual stacks to build the alleles of the same ddRAD locus. Each individual was then mapped back to the catalogue and genotyped for all the sequences using the pstacks function. The populations function was used to call SNPs present in at least 80% of the individuals per sampling site, in all the sampling sites, and with a maximum heterozygoty of 0.8 to remove potential paralogous sequences. The final filtration step was done with vcftools (*Danecek et al., 2011*) to remove SNPs with a significant departure from Hardy-Weinberg Equilibrium proportions (p-value < 0.05) in more than four sampling sites, and to remove the individuals with more than 10% missing data.

### Analyses of population structure

The filtered dataset was further thinned by removing loci with minor allele frequencies bellow 0.05 with vcftools (Danecek *et al.*, 2011), and by keeping only one SNP per ddRAD-tag using a custom R script. The final dataset consisted of 131 individuals genotyped for 3 714 SNPs with an average coverage of 52X (Figure S1) and 2.69% missing data in the overall genotype matrix. We used the R package adegenet (Jombart, 2008) to conduct a principle component analysis (PCA) and to calculate the observed heterozygosity (H_O_) per sampling site. Pairwise *F*_ST_ between sites were estimated following the method of Weir and Cockerham (1984) and their significance were tested with 1 000 permutations over loci using the R package StAMPP (Pembleton *et al.*, 2013). The effect of the IBD was tested using a Mantel test (1 000 permutations) from the R package ade4 (Dray and Dufour, 2007). The distances between a given sampling site and its closest site was obtained by measuring the kilometers along a straight line between the two sites. Since all the sites were sampled approximatively in a “stepping-stone” strategy, the distance between sampling sites separated by several sites were calculating by summing the distance to each proximal site. In order to identify linkage blocks in the dataset, we computed r^2^ (Hill and Robertson, 1968) to estimate Linkage Disequilibrium (LD) between all pairs of loci and analysed them through a network analysis with the R package LDNa (Kemppainen *et al.*, 2015). Finally, we calculated Weir and Cockerham (1984) pairwise *F*_ST_ between the two most distant sites (North Sea vs. Baltic Sea) at every single SNP to examine the variation of *F*_ST_ in the dataset.

### Analyses of demographic history

We used the two most distant sampling sites, the North Sea (NorsS) and Baltic Sea (Balt), to infer the most likely demographic history at the origin of the observed population structure across the transition zone. The dataset was filtered to retain only polymorphic SNPs in at least one of the two populations, and one SNP per ddRAD-tag to limit effects from physical LD. We used the version of δaδi software (Gutenkunst *et al.*, 2010) modified by Tine *et al.* (2014). The method uses the joint allelic frequency spectrum (JAFS) to infer the most likely population history from contrasting demographic scenarios. Without access to data from a closely related outgroup species, the inferences were performed using the folded version of the JAFS for 30 chromosomes (15 individuals) to represent the spectrum. We compared demographic models in which one ancestral population split into two derived populations for a certain period of time. In the simplest scenario, the populations diverge in total isolation (Strict Isolation: SI). Then, the possibility of exchanging migrants was added to the models by setting either a continuous period of gene flow after the population split (Isolation-with-Migration: IM), or discontinuous gene flow, following or predating an isolation phase (secondary contact: SC and Ancestral migration: AM, respectively). As the two sole populations are separated by a steep environmental gradient they are therefore likely to experience different selective pressures. Consequently, we included the indirect effect of selection in the models by adding a proportion of loci with reduced effective size to the four models, or, a proportion of loci with a reduced migration rate to the three models with gene flow (more details about the model setting can be found in Rougeux *et al*, 2018). In total, 11 simplified scenarios were compared, representing scenarios potentially involved during the colonization of the Baltic Sea. For each model, 30 independent runs were performed, from which the five best fits were retained. The goodness of fit of every model was compared using the Akaike Information Criterion (AIC). A difference bellow 10 was considered insufficient to distinguish the models (Rougeux *et al*, 2018). If several models were equivalent, the model with the fewest parameter was considered as the most likely scenario at the origin of the population split.

### Test for selection

The parameters from the best demographic scenario (Figure 2B) were used to simulate neutral divergence of two populations in msms (Ewing and Hermisson, 2010). The sampling sizes of the simulated data were equivalent to those actually sampled in the North Sea and Baltic Sea populations. From this dataset, we calculated overall expected heterozygosity (H_E_) and *F*_ST_ between populations using the program msstat (Thornton, 2003). We then used a smoothed nonparametric quantile regression to obtain the 99% and 99.9% neutral envelope of *F*_ST_ for bins of 0.02 of H_E_ from the simulated data, following the approach of Beaumont and Nichols (1996). The upper quantiles of the neutral *F*_ST_ distribution were predicted based of the H_E_ of each SNP in the observed dataset, and all the SNPs with an *F*_ST_ value above the predicted upper quantile were considered as candidate *F*_ST_ outlier loci. Finally, all RAD-sequences carrying a candidate locus for selection were aligned to the NCBI database the function blastn from ncbi-blast v.2.6.0+ (Altschul *et al.*, 1990). To finish, we used the R package HZAR to fit an allelic frequency cline for each top 0.1% outlier loci identified.

**Figure 2:**
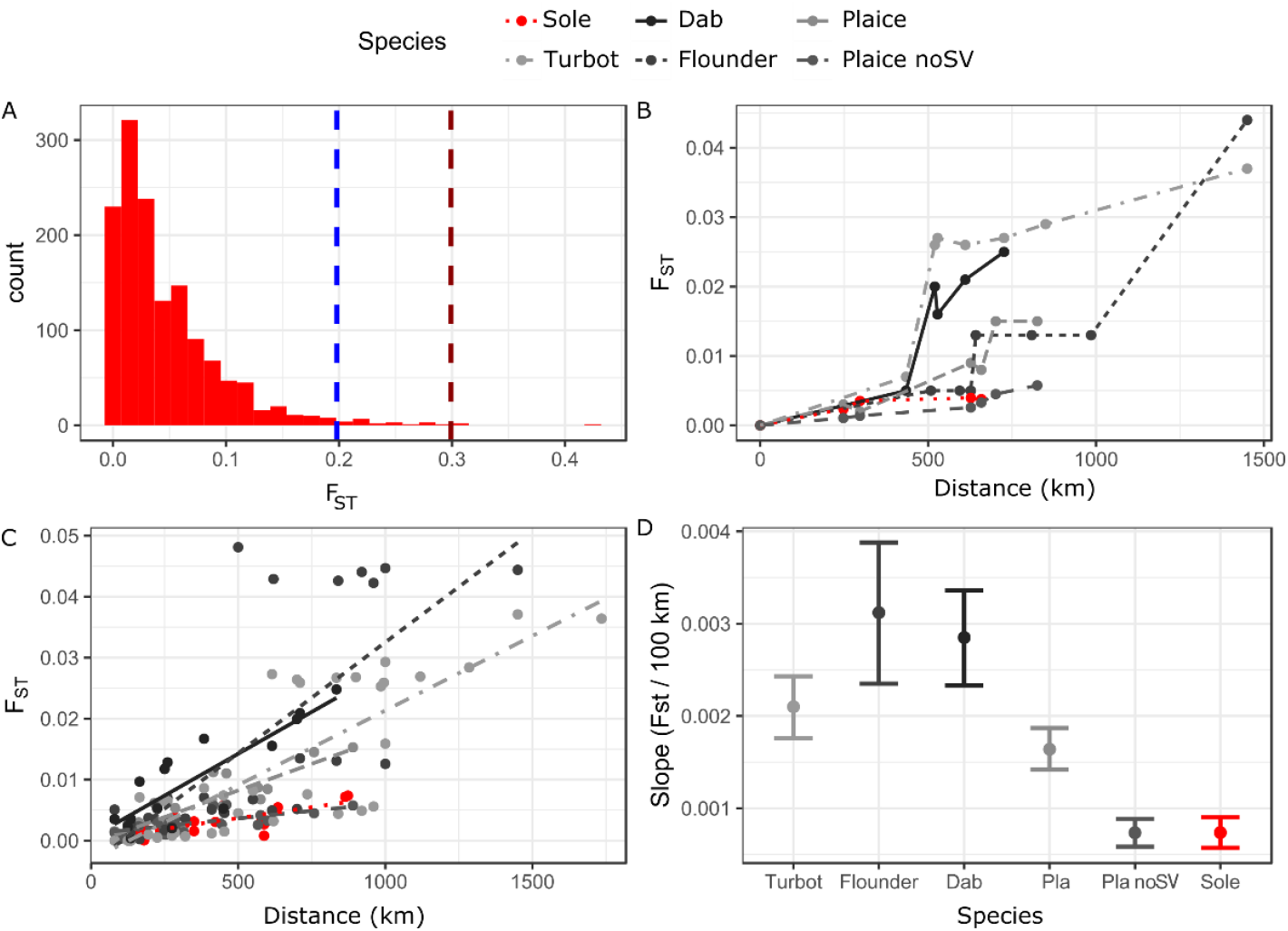
Distribution of pairwise *F*_ST_ between the North Sea and Baltic Sea across loci in sole (A), *F*_ST_ values between the North Sea and other sampling sites along the North Sea-Baltic Sea transition zone in five flatfish species (B), linear regression (C), and slope (with confidence intervals) of *F*_ST_ as a function of geographical distance (D). The vertical bars in A correspond to the upper 1 % (blue) and upper 0.1% (brown) quantiles for simulated neutral loci. The zero from the graphic B was set at a fixed point along the southern coast of the North Sea at the border between Germany and the Netherlands.

### Comparative framework

For a comparative purpose, the sole IBD pattern was compared to the IBD patterns of four other flatfish species from Le Moan *et al*. (2019a): turbot, European plaice, European flounder and common dab. Their genomic data were obtained with the same ddRAD protocol used here. For each species, we only selected the North Sea sampling sites overlapping with those sampled for sole. However, we retained all the sampling sites within the Baltic Sea, even if they were variable across species, because the sampling design for the species in the Baltic Sea accurately represents their natural distribution in the region. The plaice dataset was characterized by two large polymorphic structural variants (described in Le Moan *et al.*, 2019a,b). Consequently, the plaice IBD was analysed with and without the structural variants. For each species, we tested for IBD between pairwise *F*_ST_ and geographical distance (km) along the coast using Mantel tests, as detailed previously. Furthermore, we used linear models with sole as the focal species to compare the strength of the sole IBD relating to the IBD of the other species. These models were set by forcing the intercept to be zero. Then, we compared models with and without the interaction between species and geographical distance using the likelihood ratio test from the base package of R (Ihaka and Gentleman, 1996). The ability of the linear model to predict the genetic differences for each species was evaluated by examining the distribution of the residual variation in the best model. Finally, for each species, we extracted the estimates of the slopes between the *F*_ST_ and the geographic distance and their confidence interval to obtain a proxy for the strength of IBD.

## Results

### Sole population structure

The average heterozygosity was similar in all sampling sites (H_O_ = 0.24), except for the Baltic Sea that was marginally lower (H_O_ = 0.23). The first axis of the PCA explained 1.25 % of the total inertia and showed a relatively weak structure following the geography (Figure 1). The second axis explained 1.11% of the total inertia and was associated with individual genetic diversity (Figure 1). No apparent discrete cluster was observed with this analysis (Figure 1). Pairwise *F*_ST_ analyses confirmed weak genetic differences between sites (*F*_ST_ = 0.0026), but only two pairwise comparisons were not significantly different from 0 (Table S1). Overall, pairwise *F*_ST_ values were significantly correlated with geographical distances (r = 0.73, p = 0.007, Table 1 and Figure 2C in red). The maximum *F*_ST_ estimate of *F*_ST_ = 0.0077 was observed between the two most distant sampling sites (North Sea vs. Baltic Sea – Table S1). Pairwise *F*_ST_ values between these sites was consistently low across most of the SNPs as evidenced by the distribution of *F*_ST_ across loci (Figure 2A). Nevertheless, few markers were clearly outside the main distribution with one outlier estimate of 0.4 (five SNPs with *F*_ST_ > 0.3 on the dataset without LD filtration, Figure S2). The average LD was relatively weak (r=0.0002). Although few pairs of loci showed high values, potentially linked to physical proximity, no major linkage blocks were identified with LDna (Figure S3).

**Table 1:** Pattern of structure for the five species of flatfish along the North Sea - Baltic Sea transition zone. In order of appearance, for each species: the overall average *F*_ST_, the Mantel correlation between distance and *F*_ST_, the p-value of the correlation, the slope of the linear regression of *F*_ST_ on distance the confidence interval of the slope and the best demographic scenario at the origin of the differentiation.

| Species | Av. $F_{ST}$ | Cor. | p-value | Slope | conf. int. | Scenario |
| --- | --- | --- | --- | --- | --- | --- |
| Sole | 0.0026 | 0.73 | 0.007 | 0.0007 | [0.0006-0.0009] | IM |
| Turbot | 0.0085 | 0.82 | 0.000 | 0.0021 | [0.0018-0.0024] | SC2m (Le Moan *et al.*, 2019a) |
| Flounder | 0.0200 | 0.72 | 0.001 | 0.0031 | [0.0024-0.0039] | SC2m (Le Moan *et al.*, 2019a) |
| Dab | 0.0079 | 0.86 | 0.002 | 0.0029 | [0.0023-0.0034] | SC2m (Le Moan *et al.*, 2019a) |
| Plaice (Inv.) | 0.0048 | 0.87 | 0.001 | 0.0016 | [0.0014-0.0019] | IM2m (Le Moan *et al.*, 2019a) |
| Plaice (no Inv.) | 0.0028 | 0.68 | 0.004 | 0.0007 | [0.0006-0.0009] | IM2m (Le Moan *et al.*, 2019a) |

### Demographic history of the sole

The performance of models to fit the observed divergence was similar in the models without gene flow (SI & SI2N, AIC = 1600), but these models resulted in poor fits of the JAFS in comparison to the models with gene flow (Figure 3A and Table S2). However, all the models with gene flow performed similarly well with all the AIC values ranging between 1006 and 996. As the differences in AIC were below 10, all the models with gene flow provided equally good fits to the data (Figure 3A). Indeed, the models with discontinuous gene flow (AM and SC) converged towards isolation phases which were relatively low (Table S2) and therefore were equivalent to the model with continuous gene flow (IM). Interestingly, the model taking the possibility of selection into account did not perform better than the models without selection. Thus, we considered the IM model to be the most likely to explain population differentiation in sole (Figure 3). The inferences under this scenario provided effective size estimates of the North Sea sole population that were than the effective size of the population from the transition zone. The two populations were connected by highly asymmetric migration rates, with most of the gene flow occurring from the Baltic Sea into the North Sea (Figure 3B).

**Figure 3:**
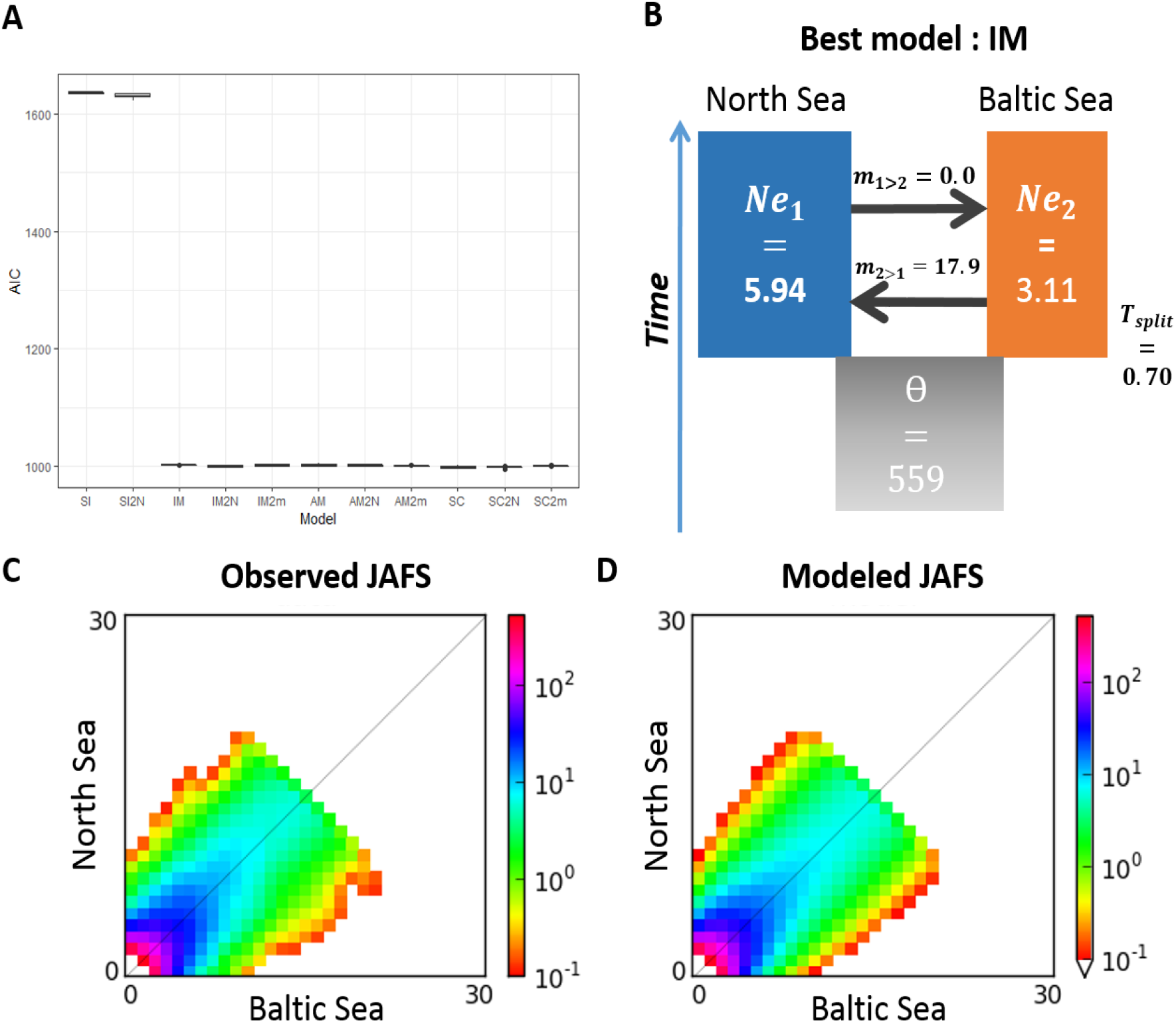
Results from the demographic inference analyses for the two populations of sole (North Sea and Baltic Sea). The AIC obtained from the five best fits of each model (A), the model IM providing the best fit with the value inferred by δaδi (given relative to theta) in B), the observed JAFS (C) and fitted JAFS from the IM model (D)

### Evidence for selection in the sole data

In total, 77 SNPs were located among the top 1% of the simulated neutral loci under the demographic scenario of Figure 3B. Only eight loci were among the upper 0.1% (Figure 4A) and most of these loci showed allele frequency clines more or less linear across the studied area, except for two loci (represented in brown and in purple in Figure 4B) which exhibited a more sudden increase/decrease in frequency. The locus shown in brown displayed the minor allele in high frequency only in the southern North Sea, while the minor allele of the locus shown in purple increased in frequency only in the three sampling sites located within the environmental gradient. Most of the outlier loci aligned to various fish genomes from the NCBI database but only four were localized in coding sequences (Supplementary file I).

**Figure 4:**
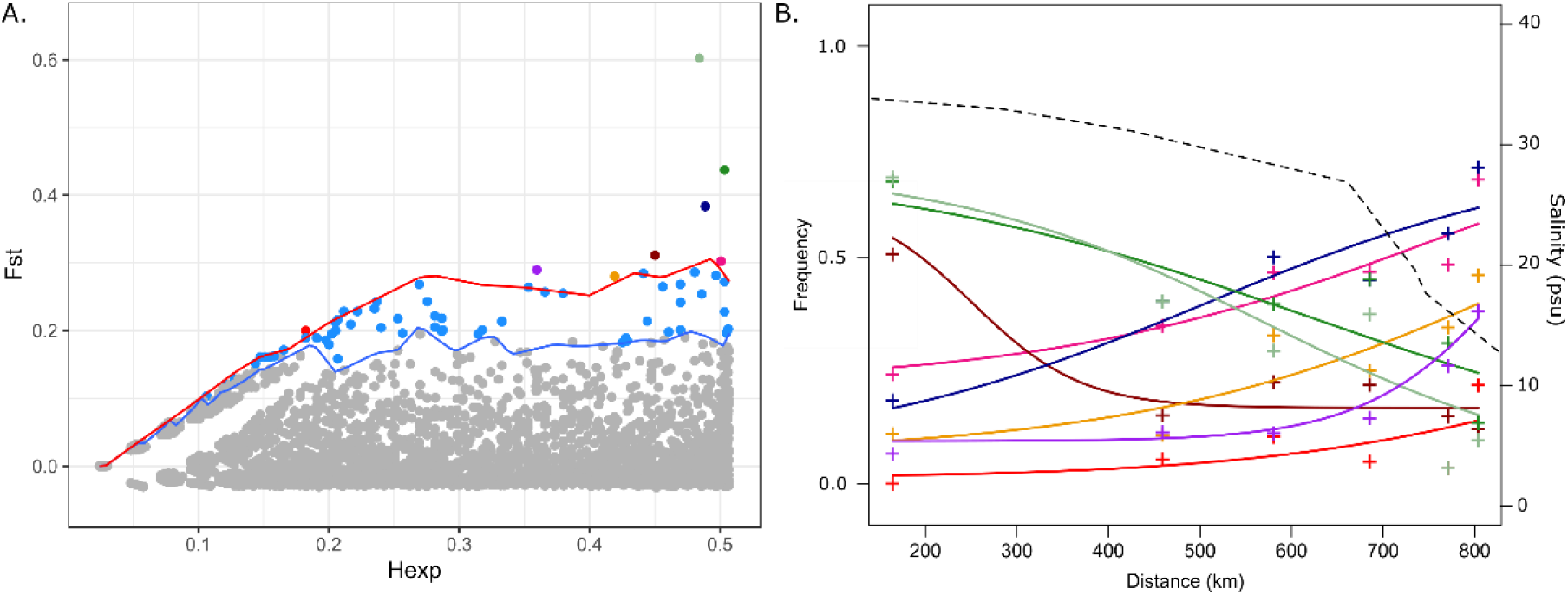
F_ST_ outlier analyses with the result from the genome scan (A), where the blue and red lines correspond to the neutral envelope of differentiation at 1% and 0.1% respectively. The envelopes were obtained from the simulation with the δaδi parameters, and the colored dots correspond to the top 1% outliers. Dots with a single individual color (other than light blue) correspond to the top 0.1%, from which the slopes of allelic frequencies are represented in B. The dashed line in B represents the salinity gradient of the transition zone. The distance are given from a fixe point along the southern coast of the North Sea at the border between Germany and the Netherlands, and each cross correspond to the allelic frequency at each sampled sites.

### Comparison of sole IBD with IBD in other flatfish species

Pairwise *F*_ST_ comparisons were significantly correlated with the geographical distance between sampling sites in all species (Table 1). The correlations ranged from r = 0.67 to r = 0.87 (Table 1). Most species, except the sole, showed clear breaks of *F*_ST_ along the transition zone (Figure 2B). In comparison, the sole displayed a weak and continuous increase of *F*_ST_ (Figure 2B-C), which consequently resulted in the lowest average *F*_ST_ of all species (Table 1). The linear models showed a significant effect of distance, species and the interaction between the two factors on the pairwise *F*_ST_ (r = 0.79, df = 5, p-value = 0.0007). This test confirmed that the strength of IBD varied across flatfish species along the transition zone. Only the plaice IBD was not significantly different from the sole IBD (Figure 2 and Table S2). The highest slopes of IBD were found for the flounder and dab, with an average increase of 0.003 units of *F*_ST_ per 100 km (Table 1). This increase was roughly four times higher than the slope inferred for sole. However, the linear models provided better predictions for sole and plaice data than for the other species, as observed in the distribution of the residuals (Figure S4). The prediction was particularly poor for the flounder dataset, with the residual not normally distributed around 0 and more associated to negative values. This is likely due to the presence of three highly structured populations of flounder in the area. Removing the putative structural variants from the plaice data resulted in patterns of IBD which were very similar to the sole IBD (Figure 2C-D).

## Discussion

We found weak population structure for sole along the North Sea – Baltic Sea transition zone. The signals of structuring were significant across most sampling sites and followed a pattern of isolation-by-distance. Inferences of demographic modelling suggest that the North Sea – Baltic Sea divergence have occurred in the face of continuous gene flow. Although few loci showed strong support for being under selection (eight loci in the 0.1% quantile), most of the genetic differentiation could be attributed to a neutral process of divergence. Therefore, this study provides an example of IBD mostly driven by neutral processes during the colonization of the Baltic Sea by a marine fish. Despite the biological similarities of the species of flatfish analyzed here, the sole showed the lowest IBD values. Only the plaice displayed an IBD pattern similar to the sole and these were the only two species that showed a similar demographic history for the colonization of the Baltic Sea basin.

### Population structure of sole

Previous studies have also reported patterns of IBD over a large geographical scale for sole, and the differences between the North Sea and the Baltic Sea were previously identified with 10 microsatellite markers (Cuveliers *et al.*, 2012) and 426 SNPs (Diopere *et al.*, 2018). However, increasing the genomic coverage in our study also revealed IBD within the transition zone, an area which was considered previously to be panmictic (Diopere *et al.*, 2018).

In previous studies, the authors hypothesized that a combination of selective and neutral processes could be driving the observed differences (Cuveliers *et al.*, 2012, Diopere *et al.*, 2018). With increasing genome coverage, we were able to model the demographic history of these populations and our results suggest that divergence was primarily driven by neutral processes. The population size of the North Sea was inferred to be two times larger than in the transition zone. This difference could have increased the effect of genetic drift in the population from the transition zone and could also explain the differences, albeit minor, of observed heterozygosity in the two populations. The low effective size in the transition zone may have resulted from a founder effect during the recent colonization of the Baltic Sea, as well as from the isolation effects in populations at the edge of species distributions (margin effect – Johannesson and Andre, 2009). Here, the isolation of the transition zone was supported by the inferred direction of gene flow, which was highly skewed towards higher migration rates from the Baltic Sea into the North Sea. Asymmetric gene flow is an important feature in fine-scale population structure in the marine environment (Riginos *et al.*, 2016). Since the North Sea is the only potential source of migrants to the Baltic Sea, this asymmetry could have reinforced the isolation of the population inhabiting the transition zone. Similar asymmetry of gene flow was found in three of the four species of flatfish studied in Le Moan *et al.* (2019a). The observed asymmetry of gene flow may be driven by the specific hydrographic conditions occurring in the transition zone. Indeed, oceanic gyres in the Skagerrak Sea limit inflow from the North Sea into the Baltic Sea and can contribute to the long-term isolation of populations in the area. Although exceptions occur, where strong inflows to the northern parts of the transition zone have been associated with mechanical mixing of cod populations (Knutsen *et al.*, 2004), other studies have not detected similarly strong effects on population mixture in other areas of the transition zone (Hemmer-Hansen *et al.*, 2019). Furthermore, local retention of juveniles has been suggested to be important for maintaining population structure in turbot occurring in the transition zone (Nielsen *et al.* 2005).

Although most of the genetic differentiation in our data was inferred to have evolved through neutral processes, we did identify a few highly differentiated outlier loci. These loci showed different clinal patterns across the transition zone, which could be a result of a disparity of selective pressures interacting with gene flow in the area. Local adaptation may act to reinforce asymmetry of gene flow if, for instance, the sole living in the transition zone are more plastic to environmental variation (Crispo and Chapman, 2010). Such plasticity would make the transition alleles, found in a wide range of salinities, more likely to migrate into the high salinity of the North Sea than putatively high, mal-adapted North Sea alleles to migrate into the less-saline transition zone. Altogether, the limited inflow to the Baltic Sea and the potential adaptation to the salinity gradient might reduce population admixture and could contribute to the more marked genetic isolation of the transition zone over a longer evolutionary time scale.

### Comparative population structure and IBD patterns

The sole populations are the least structured among all the flatfish species compared in the present study. In previous studies, it was not clear if similar processes were acting on sole population and in other species with more clear population structure in the area (Cuveliers *et al.*, 2012; Diopere *et al.*, 2018). The comparative approach applied here suggests that different processes and species-specific evolutionary histories are responsible for the diversity of IBD patterns observed in the species occurring in the transition zone. Indeed, the populations of sole and plaice, the closest in terms of ecological niches and biological traits (Diopere *et al.*, 2018), were inferred to share the same history of primary divergence (Figure 3 and Le Moan *et al.*, 2019a) and displayed similar patterns of IBD. Still, IBD in sole was substantially weaker than in plaice, although the difference was not statistically significant. The demographic models with selection performed better in the plaice but not in the sole. The differences in IBD pattern could therefore reflect different phases towards adaptation to the low salinity of the Baltic Sea, which also matches the extent of geographical distribution across the environmental gradient, where plaice is more prevalent at the lower salinities in the western parts of the Baltic Sea, while sole is confined to the western-most regions of the basin (Storr-Paulsen *et al.*, 2012). Indeed, plaice shows strong signatures of selection acting at two large structural variants (Le Moan *et al.*, 2019a,b), while our analyses of LD clusters in this study did not reveal any strong support for similar genomic signatures in sole. Removing the two chromosomes carrying the structural variants in plaice resulted in similar strength of IBD in the two species (slope = 0.0007 in both species). The patterns of structuring and IBD for sole and plaice were markedly different from those found for populations of flounder, turbot and dab (although only marginally for the latter), which were all inferred to correspond to secondary zones of hybridization (Le Moan *et al.*, 2019a). The predictive power in the linear models for the plaice and the sole were better than in the three other species (Figure S4). This suggest that the secondary contact events in the populations of dab, flounder and turbot may be responsible for the clinal structure observed in Figure 2b, which is not well captured by the linear model. Overall, the absence of isolation phases in the sole may have reduced their probability to build up genetic incompatibilities (*Bierne et al., 2011*), and thus contributed to decrease the effect of selection at linked sites likely involved in the secondary phase of hybridization in the other species (Gagnaire *et al.*, 2015), therefore leading to the relatively weak population structure observed. Once again, our results suggest that species-specific life history features play a fundamental role in their evolutionary history and in the history of the colonization of the Baltic Sea.

## Conclusions

This study provides a genomic view on the neutral processes involved in shaping the population structure of marine flatfishes along the North Sea – Baltic Sea transition zone, with particular focus on the sole. These neutral processes may be linked to a lower effective population size within the transition zone and to local hydrographic conditions leading to asymmetrical gene flow, potentially reinforced by local adaptation. The combination of these processes is thus likely acting to promote isolation of the Baltic Sea population of sole. The two species with similar ecological niches, biological traits and demographic histories (sole and plaice) showed similar patterns of isolation-by-distance. It is possible that the environmental conditions in the North Sea and the Baltic Sea could have reproduced similar neutral IBD patterns in the species sharing similar evolutionary histories and biological traits. In future work, more species should be included within a similar comparative framework to confirm if this specific trend represents a general rule associated with the differentiation of populations occurring across the transition zone and into the Baltic Sea.

## Acknowledgement

We sincerely thanks Dorte Meldrup for the preparation of the library, Henrik Baktoft and Casper Gundelund Jørgensen for their advice on the statistical tests and Romina Henriques for her comment on the manuscript. This study received financial support from The European Regional Development Fund (Interreg V-A, project “MarGen”), The European Maritime and Fisheries Fund and the Danish Fisheries Agency (project “Improvement of biological advice for common sole in Danish waters”, J. no. 33113-B-16-021)”, from Ørsted Foundation, and from the Otto Mønsteds fond.

## Supplementary Material

**Figure S1:**
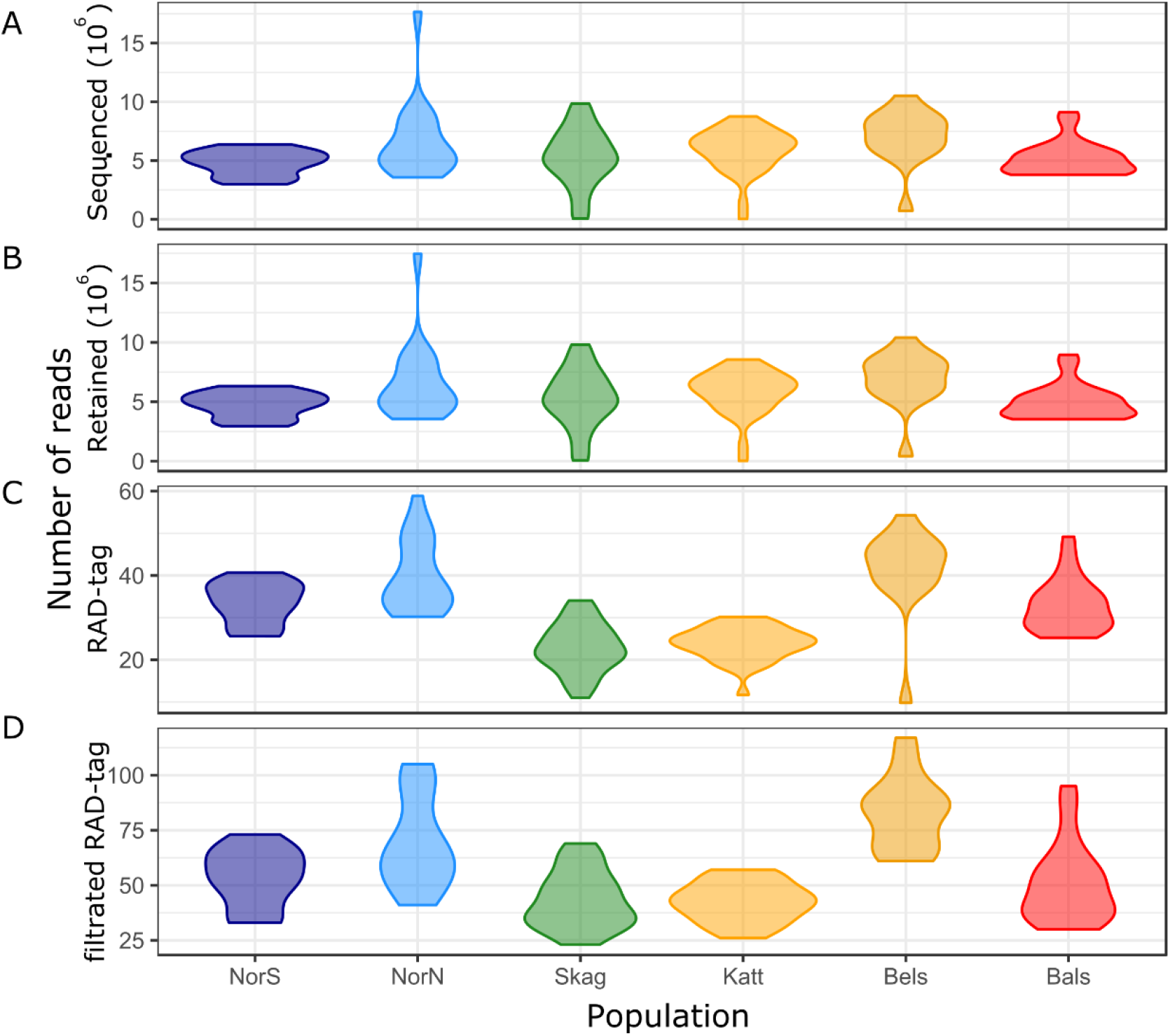
Distribution of the number of reads across individuals of sole after A. sequencing, B. retaining sequences with a phred33 quality above 10, C. stacks de-novo assembling of the RAD-tag, and D. filtration of RAD-tags

**Table S1:** Pairwise *F*_ST_ between sole populations, where values bellow the diagonal are *F*_ST_ estimates and values above the diagonal show the p-value of the permutation test. Asterisks indicate significant values above 5%

|  | Sol_4bS | Sol_4bN | Sol_20 | Sol_21 | Sol_22 | Sol_23 |
| --- | --- | --- | --- | --- | --- | --- |
| NorS | -- | 0.0000 | 0.0000 | 0.0000 | 0.0000 | 0.0000 |
| NorN | 0.0039* | -- | 0.0000 | 0.0000 | 0.0020 | 0.0000 |
| Skag | 0.0045* | 0.0023* | -- | 0.0130 | 0.1680 | 0.0030 |
| Katt | 0.0047* | 0.0035* | 0.0014* | -- | 0.0000 | 0.0000 |
| Bels | 0.0069* | 0.0040* | 0.0001 | 0.0031* | -- | 0.4920 |
| Bals | 0.0077* | 0.0038* | 0.0008* | 0.0022* | 0.0006 |  |

**Figure S2:**
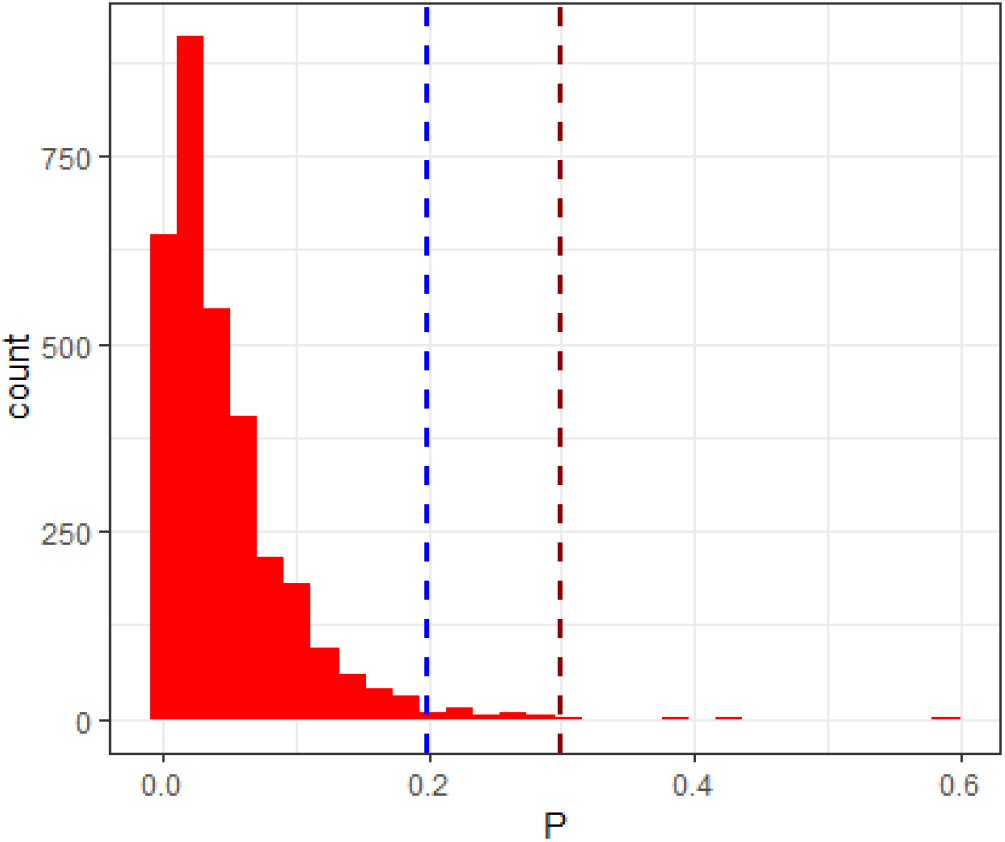
Pairwise *F*_ST_ distribution between the two most distant sampling sites for sole without pruning the data for LD

**Figure S3:**
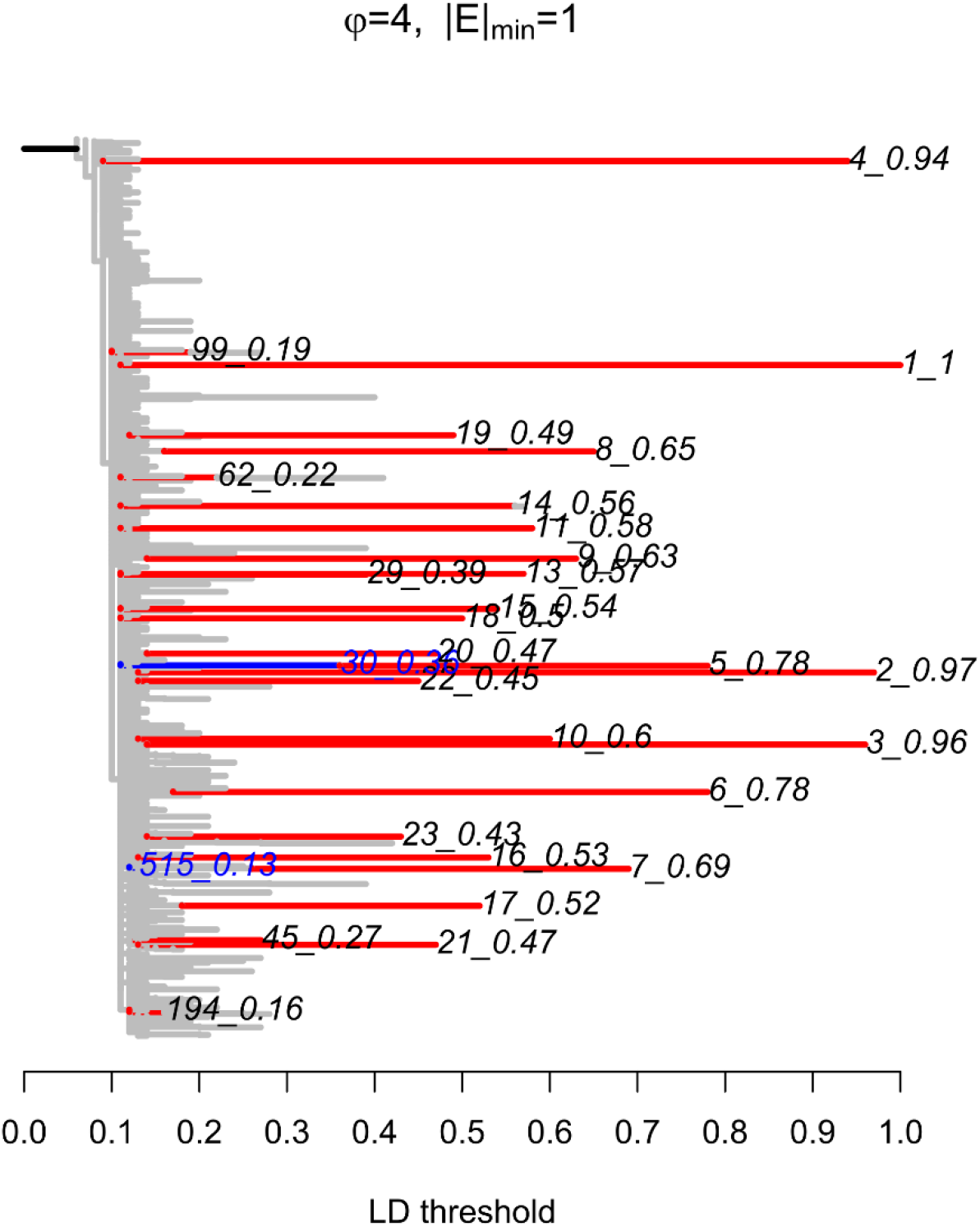
LD network between pairs of SNPs in the sole dataset. The important LD values correspond to clusters including less than four SNPs

**Table S1:** Result from the demographic inferences performed in δaδi for the best fit of each models (first columns), with, in order of appearance: the likelihood of the model (lh.); the Akaike criterion (AIC); the differences with the AIC of the best model (*Δ*_*AIC*_); the ancestral mutation rate (θ); the effective size of the North Sea and the Baltic Sea populations (N_E_1 and N_E_2, respectively); the migration rate from the North Sea into the Baltic Sea and from the Baltic Sea into the North Sea (m_1>2_ and m_2>1_); the reduced migration rate (mr_1>2_ and mr_2>1_); the time of split between the North Sea and the Baltic Sea (T_S_); the time of ancestral migration where migration is set at 0 (T_AM_); the time of secondary contact (T_SC_), and the proportion of loci with reduced migration rate / reduced effective size (P/Q)

| Model | lh. | AIC | $\Delta_{AIC}$ | $\theta$ | $N_{E1}$ | $N_{E2}$ | $rN_E$ | $m_{1>2}$ | $m_{2>1}$ | $mr_{1>2}$ | $mr_{2>1}$ | $T_s$ | $T_{AM}$ | $T_{SC}$ | P/Q |
| --- | --- | --- | --- | --- | --- | --- | --- | --- | --- | --- | --- | --- | --- | --- | --- |
| SI | -1815 | 1635 | 639 | 951 | 19,97 | 19,96 | -- | -- | -- | -- | -- | 0,17 | -- | -- | -- |
| SI2N | -1806 | 1623 | 626 | 938 | 47,34 | 55,83 | 0,19 | -- | -- | -- | -- | 0,10 | -- | -- | 0,08 |
| **IM** | **-1496** | **1002** | **6** | **560** | **5,96** | **3,11** | -- | **0,00** | **17,97** | -- | -- | **0,72** | -- | -- | -- |
| IM2m | -1491 | 998 | 2 | 553 | 6,50 | 3,14 | -- | 4,23 | 0,00 | 0,00 | 38,56 | 0,76 | -- | -- | 0,22 |
| IM2N | -1492 | 997 | 1 | 542 | 11,23 | 3,84 | 0,28 | 0,00 | 23,99 | -- | -- | 0,85 | -- | -- | 0,50 |
| AM | -1495 | 1001 | 5 | 556 | 5,88 | 6,83 | -- | 0,04 | 27,82 | -- | -- | 0,72 | 0,03 | -- | -- |
| AM2m | -1490 | 999 | 2 | 461 | 3,25 | 26,47 | -- | 21,90 | 0,00 | 1,06 | 9,54 | 1,43 | 0,00 | -- | 0,25 |
| AM2N | -1492 | 1000 | 4 | 496 | 6,61 | 11,11 | 0,25 | 17,37 | 0,17 | -- | -- | 1,09 | 0,00 | -- | 0,50 |
| SC | -1492 | 997 | 1 | 542 | 3,52 | 4,48 | 19,68 | -- | -- | -- | 1,97 | 0,67 | -- | 0,12 | -- |
| SC2m | -1490 | 999 | 3 | 550 | 6,42 | 3,30 | -- | 0,00 | 44,71 | 4,99 | 0,01 | 0,41 | -- | 0,35 | 0,74 |
| SC2N | -1490 | 996 | 0 | 502 | 10,42 | 14,52 | 0,25 | 0,00 | 13,30 | -- | -- | 0,72 | -- | 0,36 | 0,49 |

**Table S3:** Parameters estimated from the linear model between the pairwise *F*_ST_ and the geographical distance: estimates; standard-deviation error (std-error); value of the t-test (t-value), and statistical significance (p-value). Asterisks indicate IBD values significantly different from the sole above 5%

| Factor | estimates | std-error | t-value | p-value |
| --- | --- | --- | --- | --- |
| Distance | 0.000007 | 0.000008 | 0.901000 | 0.369 |
| Sole | 0.000159 | 0.003941 | 0.040000 | 0.968 |
| Dab | 0.000530 | 0.003244 | 0.163000 | 0.871 |
| Flounder | -0.003834 | 0.002424 | -1.582000 | 0.116 |
| Plaice | -0.000499 | 0.002914 | -0.171000 | 0.864 |
| Plaice noSVs | 0.001147 | 0.002914 | 0.394000 | 0.695 |
| Turbot | -0.003130 | 0.002152 | -1.454000 | 0.148 |
| Distance:Dab | 0.000020 | 0.000011 | 1.864000 | 0.065' |
| Distance:Flounder | 0.000029 | 0.000009 | 3.333000 | 0.001*** |
| Distance:Plaice | 0.000010 | 0.000011 | 0.982000 | 0.328 |
| Distance:Plaice noSVs | -0.000002 | 0.000011 | -0.199000 | 0.842 |
| Distance:Turbot | 0.000017 | 0.000008 | 2.075000 | 0.040* |

**Figure S4:**
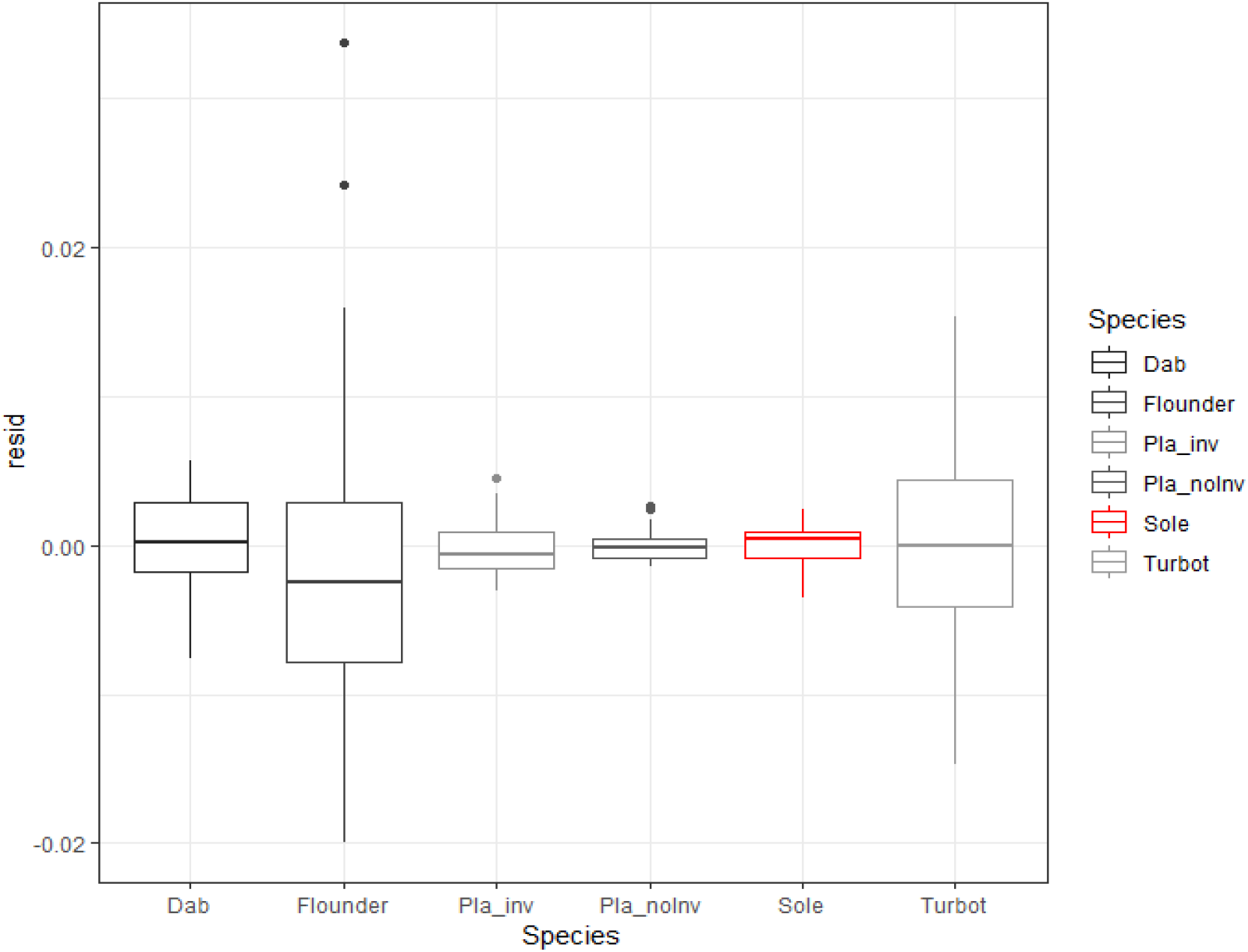
Distribution of the residuals from the linear model for each species. Lower variation indicates better fit of the linear regression

